# Enhancing visual perceptual learning using transcranial electrical stimulation: transcranial alternating current stimulation outperforms both transcranial direct current and random noise stimulation

**DOI:** 10.1101/2023.08.24.554599

**Authors:** Qing He, Xinyi Zhu, Fang Fang

**Affiliations:** School of Psychological and Cognitive Sciences and Beijing Key Laboratory of Behavior and Mental Health, Peking University, 100871 Beijing, China; Key Laboratory of Machine Perception, Ministry of Education, Peking University, 100871 Beijing, China; IDG/McGovern Institute for Brain Research, Peking University, 100871 Beijing, China; Peking-Tsinghua Center for Life Sciences, Peking University, 100871 Beijing, China

**Author notes:** These authors contributed equally (Qing He, Xinyi Zhu). Corresponding author: Fang Fang https://www.psy.pku.edu.cn/english/people/faculty/professor/fangfang/index.htm. Declaration of interests: No.

**Keywords:** transcranial electrical stimulation, tDCS, tACS, tRNS, visual perceptual learning, visual plasticity, unsupervised learning, implicit learning

## Abstract

Diverse strategies can be employed to enhance visual skills, including visual perceptual learning (VPL) and transcranial electrical stimulation (tES). Combining VPL and tES is a popular method that holds promise for significant improvements in visual acuity within a short time frame. However, there is still a lack of comprehensive evaluation regarding the effects of combining different types of tES and VPL on enhancing visual function, especially with a larger sample size. In the present study, we recruited four groups of subjects (26 subjects each) to learn an orientation discrimination task with five daily training sessions. During training, subjects’ occipital region was stimulated by one type of tES (anodal transcranial direct current stimulation (tDCS), alternating current stimulation (tACS) at 10 Hz, high-frequency random noise stimulation (tRNS), and sham tACS) while performing the training task. We found that compared with the sham stimulation, both the high-frequency tRNS and the 10 Hz tACS facilitated VPL efficiently in terms of learning rate and performance improvement, while there was little modulatory effect in the anodal tDCS condition. Remarkably, the 10 Hz tACS condition exhibited superior modulatory effects to the tRNS condition, demonstrating the strongest modulation among the most commonly used tES types for further enhancing vision when combined with VPL. Our results suggest that alpha oscillations play a vital role in VPL. Our study provides a practical guide for vision rehabilitation.

## Introduction

Our visual system is remarkably malleable (i.e., visual plasticity) throughout the entire lifespan, for both mature and degenerative brains. It has been well documented that various methods can bring about visual plasticity, such as extensive training and non-invasive brain stimulation (NIBS) techniques (He *et al*., 2022a). Regarding visual plasticity induced by extensive training, i.e., visual perceptual learning (VPL), many advancements have been made in unveiling its characteristics, brain loci, and neural manifestations, over the last three decades (He *et al*., 2022a). More importantly, VPL has been widely applied in various fields (Lu *et al*., 2016), such as low-vision rehabilitation (Huang *et al*., 2022; Levi *et al*., 1996), remediation for dyslexia (Gori *et al*., 2014; Meng et al., 2014), X-ray security screening (McCarley *et al*., 2004), sports and military training (Hadlow *et al*., 2018), and medical imaging education (Alexander *et al*., 2020). At the same time, our visual plasticity can also be induced by transcranial electrical stimulation (tES) – a non-invasive neuromodulatory technique, in which weak electrical fields are delivered through electrodes positioned on the scalp surface. There are three most common types of tES techniques categorized based on the current waveform, i.e., transcranial direct current stimulation (tDCS) (Nitsche *et al*., 2000; Priori *et al*., 1998), transcranial alternating current stimulation (tACS) (Antal *et al*., 2008), and transcranial random noise stimulation (tRNS) (Terney *et al*., 2008). All these three types of tES techniques have been found to be able to modify cortical excitability and modulate visual functions in both healthy and clinical populations (Bello *et al*., 2023), such as contrast sensitivity (Antal *et al*., 2001; Potok *et al*., 2023), visual acuity (Bocci et al., 2018; Reinhart *et al*., 2016), motion direction discrimination (Battaglini *et al*., 2023; Ghin *et al*., 2018), and peripheral target identification in a crowded environment (Battaglini et al., 2020; Chen *et al*., 2021), therefore establishing causal links between brain activity and cognitive functions (Zhang *et al*., 2019).

Currently, there is an increasing focus on modulating VPL by tES, which is both theoretically significant and practically useful. Theoretically, through selective modulation of the brain during distinct phases of learning, mechanisms that support successful task acquisition and consolidation may be more fully characterized (He *et al*., 2021; He *et al*., 2022a; He *et al*., 2022b; Wu *et al*., 2023; Yang *et al*., 2022). From the prospective of translational applications, trainees aspire to attain the greatest training benefits in the briefest timeframe. Whereas, in a typical practical-oriented application case, thousands of trials across multiple training sessions are required usually. Boosting VPL by tES is expected to accelerate learning and make subjects obtain greater benefits (He *et al*., 2022b; Herpich *et al*., 2019), which was aligned with trainees’ expectations.

Different types of tES have been adopted to modulate VPL (for a review, see He *et al*., 2022a). Concerning the effect of tDCS on modulating VPL, the results are mixed yet, and that no consistent result has been found regardless of whether tDCS was administered before task execution (offline mode) or during task execution (online mode). For instance, when tDCS was applied during task execution, anodal tDCS were found to be effective in boosting VPL in some studies (Frangou *et al*., 2018; Wu *et al*., 2022b), while in other studies no obvious modulatory effect was found (Fertonani *et al*., 2011; Herpich *et al*., 2019; Larcombe *et al*., 2018a; Larcombe *et al*., 2018b; Wu *et al*., 2022a) and even a suppressive effect was observed (Jia *et al*., 2022b). Studies using offline tDCS in VPL task showed similarly conflicting results (Pirulli et al., 2013; Pirulli et al., 2014; Wu *et al*., 2022b). By contrast, the results of modulating VPL by tRNS is relatively consistent. Specifically, high-frequency (100-640 Hz) tRNS boosts VPL effectively in both healthy and clinical populations (Camilleri *et al*., 2014; Campana *et al*., 2014; Cappelletti *et al*., 2015; Conto *et al*., 2021; Donkor *et al*., 2021). With regard to tACS, though the underlying neural mechanisms of action of tACS on the brain are relatively clear among tES techniques (Johnson *et al*., 2020; Krause *et al*., 2019; Zaehle *et al*., 2010), only limited studies have been conducted to modulate VPL. Specifically, we previously stimulated subjects’ visual cortical areas with tACS at different frequencies, we found that only occipital 10 Hz tACS was able to boost VPL and no such effect was found in other stimulation frequencies (e.g., 6, 20, and 40 Hz) (He *et al*., 2022b). Similarly, in another study, occipital 3 Hz tACS was found to show little effect on VPL (Zizlsperger *et al*., 2016). Moreover, we also found that the modulatory effect of 10 Hz tACS was absent when other cortical areas were stimulated (He *et al*., 2022b). Altogether, these studies demonstrate that tACS facilitates VPL in a frequency-and location-specific manner.

Despite advancements in VPL were observed following the application of diverse types of tES, there still lacks a systematic evaluation of the modulatory effects of tES on VPL. Notably, studies on modulating VPL by tES usually employ a relatively small sample size design, which might suffer the risk of sampling bias and limit the generalizability of those findings (Minarik *et al*., 2016). To this end, we recruited four groups of subjects to learn an orientation discrimination task, and each group received a specific form of stimulation, including sham, tDCS, high-frequency tRNS, and 10 Hz tACS during their training phase. Notably, there were approximately 30 participants in each group, which provides sufficient statistical power and representativeness to ensure the reliability and generalizability of our findings.

## Methods

### Subjects

A total of 104 right-handed healthy adults with normal or correct-to-normal vision took part in the present study. Subjects were assigned to one of four conditions (26 subjects in each condition): anodal tDCS (11 females; mean age 21.54 ± 2.42 years), high-frequency tRNS (20 females; mean age 21.77 ± 2.21 years), 10 Hz tACS (18 females; mean age 21.15 ± 4.58 years), and sham 10 Hz tACS (17 females; mean age 22.04 ± 2.89 years). The sample sizes were determined based on our previous study (He *et al*., 2022b), and all data for the 10 Hz tACS condition and the sham 10 Hz tACS condition in that study (He *et al*., 2022b) were included in the present study. A screening questionnaire was administrated for each subject before starting the study. Subjects were excluded if they met the following criteria: 1) age older than 30 years or younger than 18 years, 2) a history of neural surgery or epileptic seizures or any psychiatric or neurological disorders, 3) sleep disorders or a total sleep time less than seven hours per night over the last two weeks, 4) during ovulation phase of the menstrual cycle or pregnancy (He *et al*., 2019; He *et al*., 2022b). All experimental protocols and procedures were approved were approved by the Ethics Committee of the School of Psychological and Cognitive Sciences at Peking University. Before participation, an informed written consent was obtained from each subject.

### Apparatus and Stimulation protocol

MATLAB (R2015a, MathWorks, Natick, MA) and Psychotoolbox-3 extensions (Brainard, 1997) were used to generate and control visual stimuli, which were presented on an LCD monitor (Display + +, Cambridge Research Systems, UK) with a grey background (mean luminance = 30 cd/m^2^, width = 70 cm, spatial resolution = 1920 × 1080 pixels, refresh rate = 120 Hz). The only source of light in the room was the monitor. Subjects’ heads were fixed by a chin-and-head rest at a viewing distance of 70 cm. To ensure that subjects’ eye positions were stable within 1° from the fixation point when visual stimuli were presented, the Eyelink1000 plus eye-tracking system (SR Research Ltd., Ontario, Canada) was used to monitor subjects’ eye movement throughout the whole experiment.

The high-frequency (100–640 Hz) tRNS was delivered by a battery-powered current stimulator (DC-Stimulator PLUS; neuroConn GmbH, Ilmenau, Germany), and the current in other forms was delivered using the DC-STIMULATOR MC (neuroConn GmbH, Ilmenau, Germany), through a pair of rubber electrodes of 5×7 cm^2^. The electrodes were inserted into two soaked sponges (0.9% saline solution) and attached to the subjects’ scalp by two elastic bandages. The current intensity was constant 2.0 mA for anodal tDCS, while the peak-to-peak current intensity was 2.0 mA for tACS and high-frequency tRNS. The current in both active and sham tACS stimulation conditions was in the form of a sine wave. The phase difference between the two stimulation electrodes were set at zero. In each session of all stimulation conditions, the impedance was kept lower than 6 KΩ.

### Visual stimuli and task

The task paradigm and the stimulus set used in this study were the same as our previous studies (He *et al*., 2021; He *et al*., 2022b). Oriented Gabor patches with noise (diameter = 1.25°; spatial frequency = 3.0 cycle/°; Michelson contrast = 0.5; standard deviation of Gaussian envelope = 0.42°; random spatial phase; 25% of the pixels in these patches were replaced with random noise) were presented in the lower left quadrant of the visual field, 5° from the fixation point. In each trial, a small fixation point was presented first for 500 ms, followed by two Gabor patches with orientations of 26° and 26° + *θ* appeared 100 ms each in a random order with a 500-ms blank interval (**Fig.1**(A)). A two-alternative forced-choice (2AFC) method was used in the task, and subjects were instructed to judge the orientation change of the second Gabor patch relative to the first one (counter-clockwise or clockwise) by pressing keys. Subjects’ orientation discrimination threshold at 75% accuracy was estimated using a QUEST staircase procedure, such that the *θ* varied trial by trial (Watson *et al*., 1983). Subjects would have a rest after each staircase. No feedback was provided in all test and training sessions.

**Figure 1.**
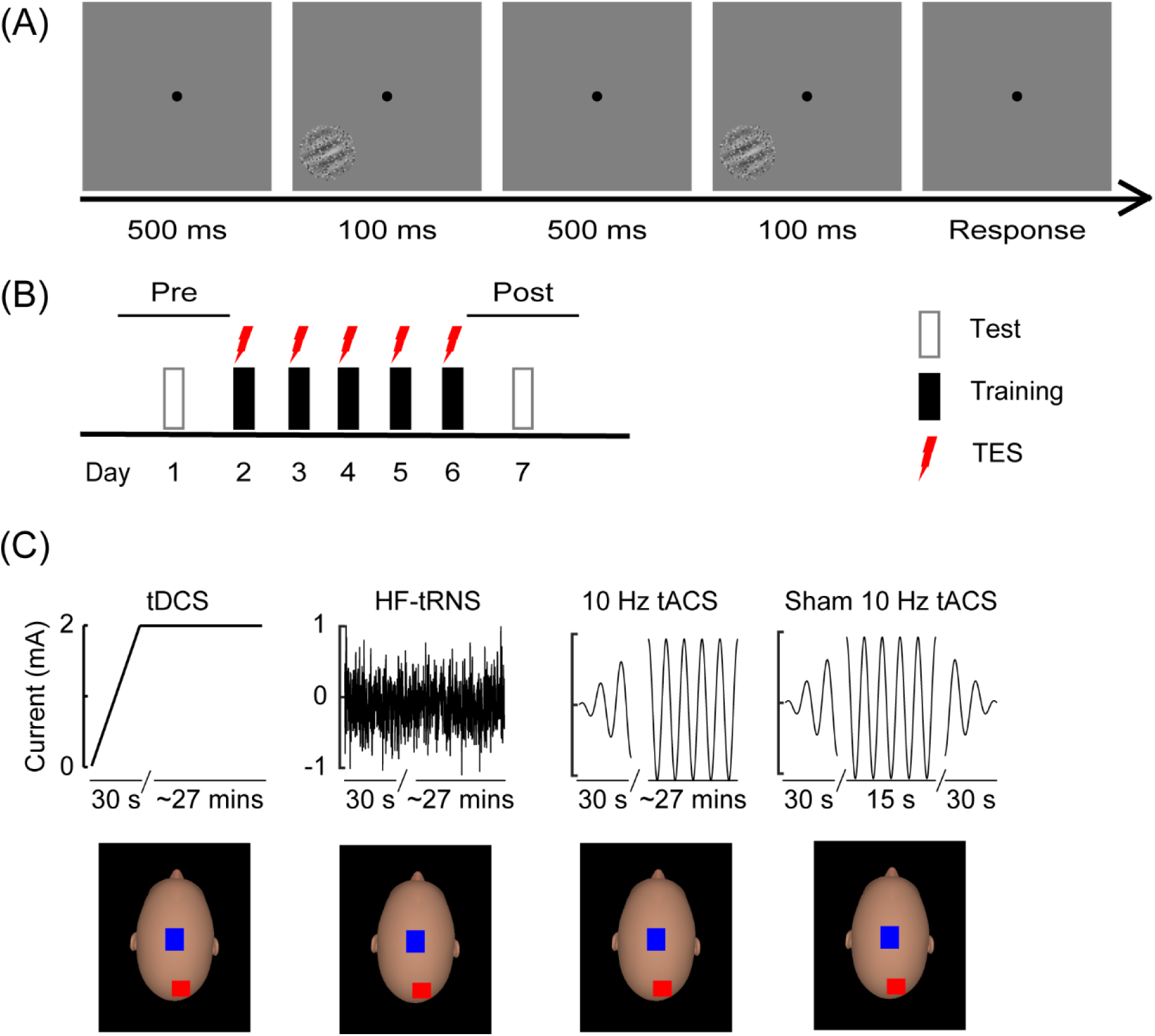
Stimuli, experimental design, and electrical stimulation protocol. (A) Schematic description of a 2AFC trial in a QUEST staircase for measuring orientation discrimination thresholds. Subjects were instructed to make a judgment of the orientation in the second interval relative to that in the first interval (clockwise or counter-clockwise), while gazing at the central fixation point. There was no feedback after each trial. (B) Experimental protocol. Subjects underwent a pre-training test, five daily training sessions, and a post-training test. The pre-training test (Pre) and post-training test (Post) took place on the days before and immediately after training, respectively. The tES was concurrently administrated during each training session. (C) Electrical stimulation protocol and montage. TES with different current waveforms (tDCS, tRNS, and tACS) were applied concurrently during each training session. Stimulation electrodes were positioned over the occipital cortex (O2) and the vertex (Cz). The electrode positions were defined by the international 10-20 EEG system. The red square and blue square denoted the anodal electrode and cathodal electrode, respectively. These head models were generated by FaceGen Modeller (version 3.4).

### Design

A single-blind, sham-controlled, between-subjects design was adopted to explore the modulatory effect of different types of tES on orientation discrimination learning. Subjects were trained on the orientation discrimination task for five consecutive days. Before and after the five daily training session, a test session was conducted, respectively, i.e., the pre-training test (Pre) and the post-training test (Post) (**Fig**.**1** (B)). Each test session and each training session consisted of six and nine QUEST staircases of 50 trials, respectively.

The hemisphere contralateral to the visual field where the visual stimuli presented was stimulated. Two electrodes were placed over subjects’ visual cortex and the vertex (i.e., O2 and Cz in the international 10-20 EEG system, respectively; **Fig**.**1**(C)). Electrical stimulation was delivered concurrently with training with a ramp-up of 30 s at the beginning of each training session.

### Statistical analysis

The threshold for each session was estimated by calculating the geometric mean of thresholds from all QUEST staircases in that session. The performance change after training was quantified by calculating the percent improvement as (*pre-training threshold* − *post-training threshold*)/*pre-training threshold* ×*100%.* All estimated thresholds were then normalized: the estimated threshold for each session was divided by the estimated threshold at Pre, then multiplied by 100%. In order to describe the process of threshold change during the learning course, we fitted the learning curves of normalized orientation discrimination threshold across all sessions using a power function:

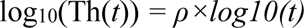

In this function, Th represents the predicted normalized threshold, *t* is the number of training sessions, and *ρ* is the learning rate (Yang *et al*., 2020). The threshold declines with training, such that the value of ρ should be negative, and a smaller value indicates a faster learning speed. To minimize the sum of squared differences between model predictions and observed values, a nonlinear least-square method was implemented in MATLAB.

A mixed-design analysis of variance (ANOVA) using *Condition* as a between-subjects factor (tDCS, tRNS, tACS, and sham) was used to analyze the orientation discrimination thresholds. Raw thresholds at Pre, learning rates and percent improvements were analyzed by ANOVA using *Condition* as a between-subjects factor. We used the Benjamini-Hochberg method (BH) to control false discovery rate (FDR) for multiple comparisons. Partial eta squared (<inline) and Cohen’s *d* were calculated to measure the effect size for ANOVAs and *t*-tests. Statistical analyses were conducted using R (R Core Team, 2023).

## Results

### Training improved task performance substantially

First, a one-way ANOVA with *Condition* (tDCS, tRNS, tACS, and sham) as a between-subjects factor on the thresholds at Pre was conducted to examine the baseline variance between all enrolled conditions. The statistical results showed that the main effect of *Condition* was not significant (*F*_(3, 100)_ *=* 0.37*, p =* 0.79, 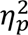 *=* 0.01), demonstrating that subjects had comparable baseline performance across all stimulation conditions. Next, to assess the learning effect on the trained task, we employed a mixed-design ANOVA that incorporated *Session* (Pre and Post) as a within-subjects factor, *Condition* as a between-subjects factor, and subjects’ orientation discrimination thresholds as the dependent variable. The statistical results showed that the main effect of *Session* (*F*_(1, 100)_ = 207.89, *p* < 0.001, 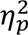 = 0.68) and the interaction between *Session* and *Condition* (*F*_(3, 100)_ = 3.29, *p* = 0.02, 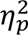 = 0.09) were significant, but the main effect of *Condition* (*F*_(3, 100)_ = 0.95, *p* = 0.42) was not significant. Further analyses showed that the thresholds at Post were lower than those at Pre for all stimulation conditions (sham: *t*(25) = 4.33, *p*_adj_ < 0.001, Cohen’s *d* = 0.85; tDCS: *t*(25) = 6.77, *p*_adj_ < 0.001; *Cohen’s d* = 1.33; tRNS: *t*(25) = 10.21, *p*_adj_ < 0.001, *Cohen’s d* = 2.00; tACS: *t*(25) = 9.01, *p*_adj_ < 0.001, *Cohen’s d* =1.77). Importantly, no feedback was provided during the training course, suggesting that subjects acquired the improved ability to discriminate the trained stimuli in an unsupervised learning manner.

**Figure 2.**
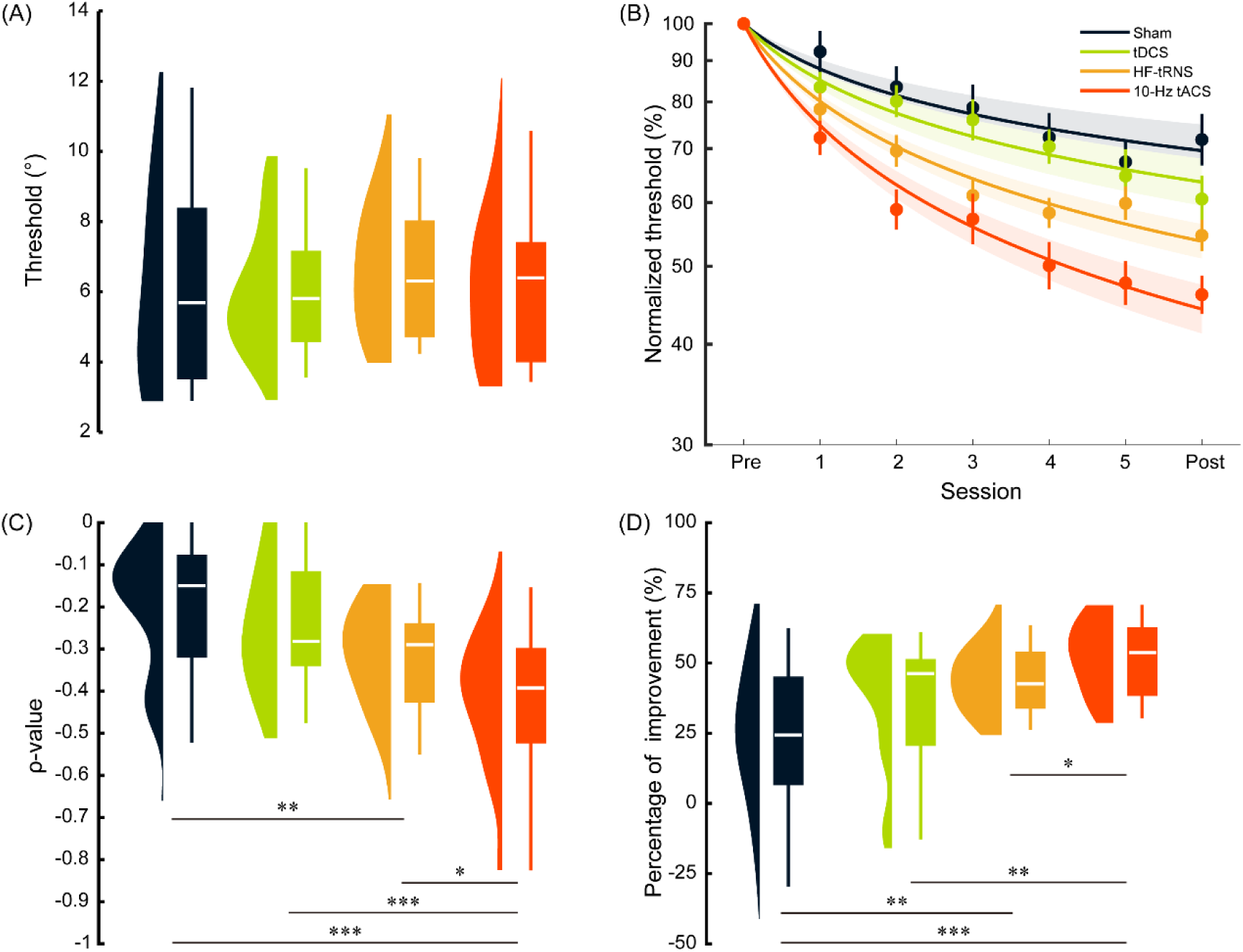
Main results. (A) Thresholds at Pre. (B) Normalized learning curves. Dots represent averaged thresholds across subjects at different test and training sessions, and lines represent fitted learning curves using a power function. Note: the y-axis is displayed on a logarithmic scale. (C) Learning rate for each condition, a smaller value indicates a faster learning speed. (D) Percentage of improvement in orientation discrimination performance. \**p* ≤ 0.05; \*\**p* ≤ 0.01; \*\*\**p* ≤ 0.001. Error bars denote 1 SEM (standard error of the mean) across subjects

### Learning rate was modulated by tES

The efficiency of performance improvement in learning is typically quantified by learning rate. We examined learning curve differences between different stimulation conditions. A one-way ANOVA revealed that the main effect of *Condition* on learning rate (ρ) was significant (*F*_(3,100)_ = 9.20, *p* < 0.001, 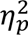 = 0.22), demonstrating that subjects in different stimulation conditions showed different learning rates. Specifically, pairwise *t*-tests showed that there was no significant difference between the sham condition and the tDCS condition in learning rate (*t*(50) = 0.90, *p*_adj_ = 0.38, *Cohen’s d* = 0.25), while the learning rates in both the tRNS and the tACS conditions were significantly greater than that in the sham condition (tRNS vs. sham: *t*(50) = 2.90, *p*_adj_ = 0.01, *Cohen’s d* = 0.80; tACS vs. sham: *t*(50) = 4.44, *p*_adj_ < 0.001, *Cohen’s d* = 1.23). Compared with the tDCS condition, the learning rate in the tACS condition was higher (*t*(50) = 3.76, *p*_adj_ = 0.001, *Cohen’s d* = 1.04), but no such effect was found in the tRNS condition (*t*(50) = 2.03, *p*_adj_ = 0.06, *Cohen’s d* = 0.56). Moreover, the learning rate in the tACS condition was higher than that in the tRNS condition (*t*(50) = 2.27, *p*_adj_ = 0.04, *Cohen’s d* = 0.63). In short, the application of 10 Hz tACS during training yielded the most rapid acceleration of the orientation discrimination learning.

### Performance improvement was modulated by tES

The space for improvement in learning is of great concern to both basic and clinical researchers. Here, we examined performance improvement differences between different stimulation conditions. Performance improvement between stimulation conditions was significant, which was revealed by a one-way ANOVA (*F*_(3,100)_ = 9.69, *p* < 0.001, 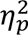 = 0.23). Further analyses showed that the percent improvement in the 10 Hz tACS condition was significantly higher than those in all other stimulation conditions (tACS vs. sham: *t*(50) = 4.85, *p*_adj_ < 0.001, *Cohen’s d* = 1.35; tACS vs. tDCS: *t*(50) = 3.08, *p*_adj_ = 0.008, *Cohen’s d* = 0.85; tACS vs. tRNS: *t*(50) = 2.33, *p*_adj_ = 0.04, *Cohen’s d* = 0.65). Notably, the performance improvement in the tRNS condition was significantly higher than that in the sham condition (*t*(50) = 3.55, *p*_adj_ = 0.003, *Cohen’s d* = 0.98). All other differences among the stimulation conditions were not significant (tDCS vs. sham: *t*(50) = 1.74, *p*_adj_ = 0.11, *Cohen’s d* = 0.48; tDCS vs. tRNS: *t*(50) = 1.60, *p*_adj_ = 0.12, *Cohen’s d* = 0.44). In short, after training with concurrent occipital 10 Hz tACS, subjects exhibited the most pronounced improvement in their performance on the trained orientation discrimination task.

In summary, subjects’ task performance improved with training from comparable initial performance levels. While the modulatory effects of tES on VPL were strongly dependent on the stimulation type. Specifically, anodal tDCS applied during executing the orientation discrimination task had little effect on modulating the acquisition of learning to discriminate the stimuli. By contrast, both high-frequency tRNS and 10 Hz tACS were capable of facilitating orientation discrimination learning, in terms of both learning rate and overall performance improvement achieved. Finally, occipital 10 Hz tACS showed the best modulatory effects.

## Discussion

In the present study, we evaluated the effect of several most common types of tES techniques (i.e., tDCS, tRNS, and 10 Hz tACS) on the further enhancement of vision by modulating VPL. Our results revealed distinct modulatory effects of tES with different current forms when applied during learning on the orientation discrimination task. Specifically, compared with the sham stimulation, anodal tDCS showed little effect on modulating VPL, while both high-frequency tRNS and 10 Hz tACS were effective in boosting VPL. Additionally, 10 Hz tACS exhibited a superior modulatory effect compared to the other current stimulation forms. To the best of our knowledge, the current study is the first to systematically evaluate the effects of various types of tES techniques on modulating VPL. Importantly, compared to all previous studies investigating the modulatory effects of tES on VPL, the present study has the largest sample size (∼30 subjects per condition), and thus ensuring sufficient statistical power for between-condition comparisons. In short, our findings will provide guidance for translational applications of combing VPL and tES, and will also provide insights into neural mechanisms of VPL.

We found that anodal tDCS applied concurrently with training had no effect on modulating VPL, which is consistent with previous studies in short-term orientation discrimination learning (Fertonani *et al*., 2011), or in multi-session learning on motion direction discrimination (Fertonani *et al*., 2019; Herpich *et al*., 2019; Larcombe *et al*., 2018a; Larcombe *et al*., 2018b; Wu *et al*., 2022a). However, our results are not consistent with some studies in which anodal tDCS was effective in facilitating VPL (Olma *et al*., 2013; Sczesny-Kaiser *et al*., 2016; Van Meel *et al*., 2016; Wu *et al*., 2022b), or even impairing VPL with a short training session (Grasso *et al*., 2021; Grasso *et al*., 2020; Jia *et al*., 2022a; Learmonth *et al*., 2015). The discrepancy between these studies mentioned above can be caused by many factors, such as training tasks, training regimes, and stimulation settings. Of note, a small sample size was adopted in previous studies, which may bias the results (Minarik *et al*., 2016). Here, using a relatively large sample size design, we found that applying anodal tDCS concurrently during training had a negligible effect on enhancing VPL. The null effect of anodal tDCS on modulating VPL during training might be caused by the timing of stimulation applied. Previous studies found that anodal tDCS applied prior to task execution (i.e., the offline mode) was effective in boosting VPL (Pirulli *et al*., 2013), while no such modulatory effect was found when anodal tDCS was administrated during perform the orientation discrimination task (i.e., the online mode) (Pirulli *et al*., 2013), just like the finding in the present study.

We found that high-frequency tRNS was effective in boosting VPL, which is consistent with previous studies (Camilleri *et al*., 2016; Camilleri *et al*., 2014; Contemori *et al*., 2019; Conto *et al*., 2021; Herpich *et al*., 2019). The consistent modulatory effects of high-frequency tRNS on VPL might be a result of strengthened attentional network. Functional connectivity within the attentional network was found to be increased after high-frequency tRNS, which was positively correlated with the magnitude of improvement in task performance (Conto *et al*., 2021). Additionally, application of high-frequency tRNS led to a decrease in blood oxygenation level dependent (BOLD) responses during task execution (Chaieb *et al*., 2009; Saiote *et al*., 2013), and reduced hemodynamic response (e.g., the amplitude of HbO2 and HbT responses) (Snowball et al., 2013). Considering that the BOLD responses are negatively correlated with alpha/beta band power typically (Bastos *et al*., 2015; Pang *et al*., 2018; Scheeringa *et al*., 2016), which is highly correlated with spatial attention (Clayton *et al*., 2015; Kelly *et al*., 2006), it is reasonable to speculate that high-frequency tRNS applied during training could increase alpha/beta band power and consequently boost learning by strengthening top-down modulation (Clayton *et al*., 2015; Conto *et al*., 2023).

We found that occipital 10 Hz tACS boosted VPL efficiently, which was consistent with our previous study (He *et al*., 2022b). In our previous study, we found that tACS boosts orientation discrimination learning in a frequency and location-specific manner. Specifically, occipital 10 Hz tACS administrated during training made subjects learn to discriminate the orientations of task stimuli faster and achieve a greater improvement, compared with the sham 10 Hz stimulation condition; further, the modulatory effect was absent in other stimulation conditions, such as occipital tACS at other alternating frequencies (6, 20, and 40 Hz) and 10 Hz tACS over other cortical regions (He *et al*., 2022b). Additionally, in another study, researchers found that tACS at 3 Hz was not effective in modulating VPL (Zizlsperger *et al*., 2016). Taken together, these results demonstrate that occipital tACS modulates VPL in a frequency-specific manner, providing strong evidence for the vital role of alpha oscillations in gating VPL (Bays *et al*., 2015; Michael *et al*., 2023).

Further, we also found that occipital 10 Hz tACS showed a stronger modulatory effect than occipital tRNS. It should be pointed out that this study is the first investigation to compare the modulatory effects of tACS and tRNS on VPL. Differences found in the modulatory effects on VPL between 10 Hz tACS and tRNS might reflect the inherent dissimilarities between these two types of tES techniques in modulating attention-related cognitive processing. It has been clearly demonstrated that 10 Hz tACS can directly entrain alpha power by aligned phase coherence or spike-timing (Huang *et al*., 2021; Johnson *et al*., 2020; Krause, *et al*., 2019), such that attention-related cognitive processing is strengthened by occipital tACS at 10 Hz directly. In contrast, high-frequency tRNS may modulate attentional processing in a relatively indirect manner, such as by attenuating BOLD response. The superior modulatory effects of occipital 10 Hz tACS on VPL compared to high-frequency tRNS may be attributed to different efficacies to mediate attentional processes. Here, this explanation is called the **efficacy of attention hypothesis**. Alternatively, we speculate that the power produced by these stimulation techniques cause the observed modulatory effects here. We propose that VPL can only be modulated by external electrical fields within a certain power range. If the power of a given stimulation falls outside of this range, it will not be effective in modulating VPL. Conversely, when the power is optimized, it produces the best modulatory effects. In this context, both 10 Hz tACS and high-frequency tRNS generate power within the range capable of modulating VPL, but the power generated by 10 Hz tACS is closer to the optimal range, and thus the former is more efficient in modulating VPL. This explanation is called the **power of stimulation hypothesis**. It needs to be emphasized that both the efficacy of attention hypothesis and the power of stimulation hypothesis need to be examined in the future.

Our findings will provide a practical guide for vision rehabilitation. VPL has been widely adopted to restore low-vision in neuro-ophthalmology (Lu *et al*., 2016). In practice, it is a common desire of patients and their families to achieve the best effect in a short course of treatment (He *et al*., 2022a). However, in a typical clinic-oriented study, weeks to months of training are required, consisting of thousands of trials, which is a significant economic and psychological burden on both the patients’ family and the wider society (Herpich *et al*., 2019). Therefore, developing methods that can speed up the process and improve the treatment effectiveness is currently a hot research topic. Our results showed that 10 Hz tACS over visual areas during training accelerated the learning process and maximized learning gains in healthy adults, which is expected to be significant in patients who need to learn or relearn skills due to injuries or illnesses. Overall, our findings have the potential to significantly impact the field of vision habilitation (Wu *et al*., 2023), and further studies will help to fully understand the effectiveness of this method for patients.

## Acknowledgements

This work was supported by the National Science and Technology Innovation 2030 Major Program (2022ZD0204802) and National Natural Science Foundation of China (31930053). Qing He was supported in part by the Postdoctoral Fellowship of Peking-Tsinghua Center for Life Sciences.

